# Avian Atlas: Unveiling the Diversity Divide in Desert Realms between Tropics

**DOI:** 10.1101/2024.05.28.596345

**Authors:** Manasi Mukherjee, Mitali Mukerji

## Abstract

Understanding diversity gradients across global deserts remains a significant challenge. The unique characteristics of deserts complicate conservation efforts for these biomes, which are highly affected by climate change. Here, we conduct a comprehensive study of ecological traits to determine the behavior and uniqueness of avian communities in ten major tropical deserts, using crowd-sourced avian diversity data from GBIF.org as a key analytical tool. Our results reveal significant variations in bird diversity among tropical deserts, with heightened diversity in deserts near the Tropic of Cancer (TCan) compared to those near the Tropic of Capricorn (TCap).

The analysis of ecological traits indicates that, unlike TCap deserts, TCan deserts have a higher prevalence of migratory species. This is facilitated by a broader niche breadth among sedentary bird species, which reduces niche competition and allows the influx of migratory invertivores. This study is the first to identify a differential avian diversity gradient within tropical deserts, demonstrating that avian species richness in deserts is more closely linked to tropical locations than to realm classifications.

Recognizing the vulnerability of TCap deserts and the diverse trophic roles played by avian species, our analysis underscores the need for targeted conservation strategies. Protecting the unique avian diversity in TCan deserts and mitigating extinction risks in TCap deserts are essential steps to ensure the resilience and sustainability of these critical ecosystems.

## 1. Introduction

Deserts—the second largest terrestrial biome on Earth exhibit surprisingly limited yet unique biodiversity (Mace et al., 2005; Ward, 2010). This challenges the species-area theory, which posits that larger areas should support more species. This paradox has been explored through studies focused on specific aspects such as desert biodiversity, functional traits, latitudinal gradients Gaston (2000), and geographic climate (Coelho et al., 2023), often conducted in isolation. These studies (Sivaperuman and Baqri, 2013; Wang et al., 2017; Alatawi et al., 2018; Conradie et al., 2019; Iknayan and Beissinger, 2020; Fei et al., 2022; Mukherjee et al., 2023) have reinforced the view that extreme conditions are the primary drivers of limited diversity in deserts. Consequently, there remains a significant gap in understanding the large-scale functioning of deserts globally, particularly in discriminating their unique characteristics and regional differences.

In our previous work, we have demonstrated the use of crowd-sourced bird diversity data in delineating the major ecoregions within a desert (Mukherjee et al., 2023). Such biogeographic distribution patterns in birds are known to be correlated to the trophic niche breadth (Fargallo et al., 2022). However, our further investigation on geographic distinctness of desert through mapping avian biodiversity with observational data, revealed that trophic niches not only play a crucial role in determining regional diversity but also the influx of migratory species (Mukherjee et al., 2024). These migratory birds due to their wide distribution (Pereira and Cooper, 2006), sensitivity to environmental changes (Smits and Fernie, 2013) and extensive existing data (Fraixedas et al., 2020), are often chosen as indicators for studying large-scale ecosystem functioning.

In this study, we extend our previous research to a global scale by utilizing crowd-sourced eBird data on avian biodiversity (2,374 bird species) from the Global Biodiversity Information Facility (GBIF.org, 2024 comprehensive global avian ecological data from Tobias et al. (2022)). Retrieving precise ecological information that maps to specific ecoregions is a challenging task. For the first time, we used an innovative approach to accurately extract avian biodiversity data specifically from desert regions using the GBIF polygon tool. The objectives of this study were to: (a) determine if avian diversity is consistent across deserts globally, (b) understand if functional traits, such as trophic niches, reveal unique characteristics of deserts, and (c) identify if variations in trophic niche structures influence migratory bird patterns across deserts. This is the first of its kind of attempt to investigate the behavior and uniqueness of ten major tropical deserts using avian diversity as a key analytical tool.

This work highlights the significant variations in bird species richness, migratory category and trophic niches among deserts of Tropic of Cancer (henceforth noted as TCan) and Tropic of Capricorn (henceforth noted as TCap). It also provides a scientific basis for identifying characteristic differences among global deserts for targeted conservation strategies and the need to protect the unique avian diversity in TC deserts as well as mitigate extinction risks in TP deserts.

## 2. Methods

### 2.1. Data collection

For the present investigation, a comprehensive selection encompassing ten major desert biomes dispersed across five distinct continental regions was undertaken to study the differences in avian diversity among desert biomes. These deserts include the Sahara and Kalahari in Africa; the Arabian, Thar, and Gobi deserts in Asia; the Great Basin and Chihuahuan deserts in North America; the Atacama and Patagonian deserts in South America; and the extensive Great Australian Desert, which comprises the Great Victoria, Sandy, Tanami, Simpson, Gibson, Strzelecki, and Sturt Stony deserts in Australia (Fig. 1).

**Figure 1:**
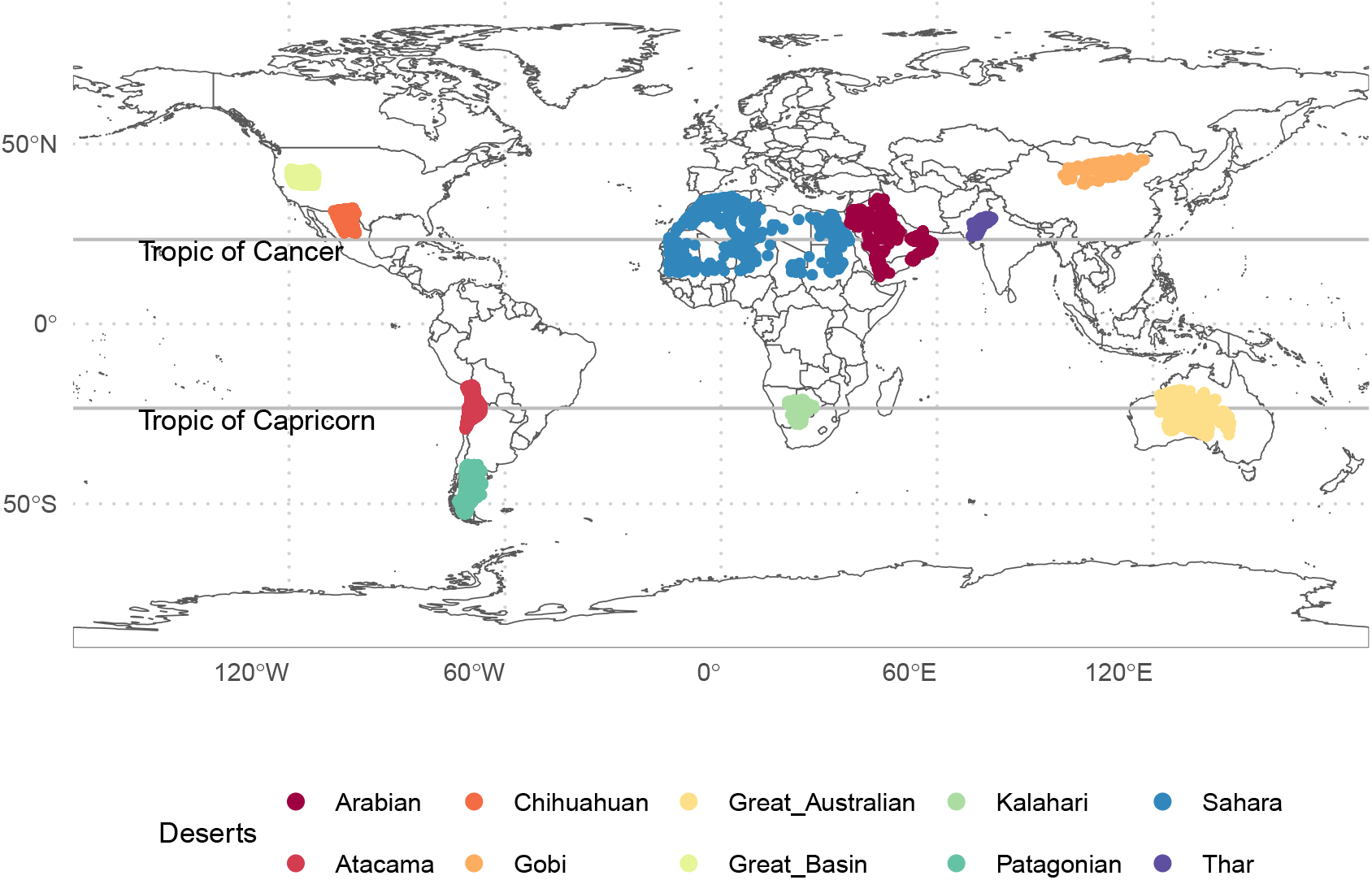
Global distribution of avian species richness in ten major deserts, as depicted on the map. Observation locations are marked with dots, which are positioned according to the latitude and longitude data sourced from eBird contributions. A higher density of dots reflects a greater volume of avian observations at those locations. This spatial analysis encompasses avian species richness data curated from the Global Biodiversity Information Facility (GBIF.org, 2024) for these selected desert ecosystems. Among the studied deserts, the Sahara, Arabian, Thar, Gobi, Chihuahuan, and Great Basin deserts are situated within the Tropic of Cancer, while the Great Australian, Kalahari, Atacama, and Patagonian deserts are located within the Tropic of Capricorn.

Avian biodiversity data, specifically species richness, pertaining to these ten selected desert ecosystems were curated from the Global Biodiversity Information Facility (GBIF.org, 2024). Given that the avian datasets in GBIF are primarily aggregated through eBird and are not explicitly categorized by desert biomes, a methodological approach. We selected the location tab from the occurrence (main menu) bar of the website. Using the coordinates from (Jung et al., 2020), the polygonal representations that closely approximate the geographical boundaries of the selected deserts was employed to curate the avain species richness data.

The variance in correlation patterns between avian behavioral traits across deserts located within the TCan and the TCap were analysed employing data from AVONET (Tobias et al., 2022). The AVONET dataset categorizes bird species into following migratory behaviour:

- Migratory: i.e. majority of population undertakes long-distance migration
- Partial Migratory: i.e. minority of population migrates long distances, or most of population undergoes short-distance migration, nomadic movements, distinct altitudinal migration, etc.
- Sedentary

### 2.2. Data Analysis

The dataset was organized to represent the occurrence and non-occurrence of various avian species across different desert environments. This involved positioning the bird species as columns and the deserts as rows. Within this arrangement, a binary system was adopted for each cell: a ‘1’denoted the presence of a bird species within a desert, while a ‘0’indicated its absence. Consequently, this structured approach yielded a matrix, denoted as *M*, characterized by binary values reflecting the presence or absence of bird species across various deserts. The entire program was implemented using ‘R’, a widely recognized open-source statistical software.

#### 2.2.1. Ecological Cluster Identification

The initial step involved determining the similarities among deserts regarding their avian species richness. To visualize these similarities, multidimensional scaling (MDS), a non-linear technique for dimensionality reduction (Tao Shen, 2009), was employed. By applying MDS to matrix *M* , it was possible to depict each desert within a two-dimensional space. Following this, the k-means clustering algorithm (MacQueen, 1967) using Euclidean distance was implemented to categorize deserts into subgroups that exhibited similarities in terms of avian species richness. The optimal number of clusters, *k*, was determined through the Elbow method (Thorndike, 1953), which involved computing and plotting the total within-cluster sum of squares (*wss*) against various *k* values (Fig. 2). It was observed that increasing *k* led to a reduction in *wss*; however, beyond *k* = 3, the reduction in *wss* became markedly less significant. As a result, *k* = 3 was selected as the appropriate number of clusters for our analysis.

**Figure 2:**
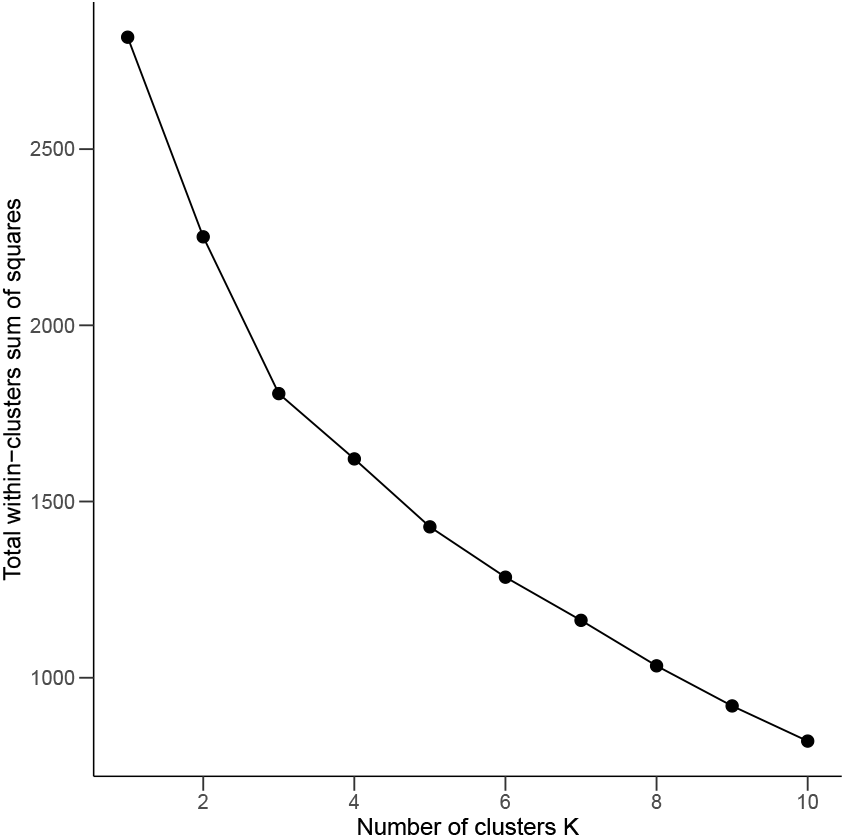

#### 2.2.2. Statistical analysis

Following the identification of desert clusters via Multidimensional Scaling (MDS) and k-means clustering, a correlation plot (corrplot) was implemented to explore and quantify the degree of similarity and statistical significance among the deserts of these clusters. Each cell in the corrplot corresponds to the correlation coefficient between two deserts, providing insight into how closely related they are in terms of the variables considered.

A two-way Analysis of Variance (ANOVA) was employed to assess the impact of migratory behavior and geographic location on avian species counts. The ANOVA was structured to evaluate the main effects of two factors: migratory category (Migratory, Partial Migratory, Sedentary) and tropical zone (TCan and TCap) as well as their interaction. This statistical method enabled us to determine whether differences in avian counts were statistically significant across the three migratory categories and between the two distinct tropical zones.

Following the identification of significant effects from the ANOVA, we conducted Tukey’s Honest Significant Difference (HSD) test as a post-hoc analysis to pinpoint specific differences between group means. This test provided a detailed comparison between each pair of migratory categories within each tropical zone, allowing for a nuanced interpretation of how migratory behaviors influence avian biodiversity across different geographic locales.

#### 2.2.3. Migratory and Trophic niche difference analysis

The AVONET dataset, which aggregates morphological, ecological, and geographical attributes for 10,662 avian species as documented by Tobias et al., (2022), served as the primary data source for analyzing variations in avian migration patterns across distinct desert aggregations. Within this comprehensive dataset, a subset of 2,374 bird species identified from the selected deserts was cross-referenced, revealing a concordance with 1,930 species in the AVONET dataset. The remaining species were excluded from further migratory and trophic niche analyses due to the lack of matching data. The dataset delineates avian migration status into three categories: Migratory (M), Partially Migratory (PM), and Sedentary (S). The study aimed to elucidate potential differences in migratory behaviors among birds inhabiting the delineated desert groups. The classification of trophic niches within this dataset encompasses a diverse range of feeding behaviors, including Aquatic Predator, Frugivore, Granivore, Herbivore (Aquatic), Invertivore, Nectarivore, Omnivore, Scavenger, and Vertivore.

Bird species representative of each desert were categorized according to their migratory status. Subsequently, a percentage stacked bar graph was constructed to visually depict the variation in migratory composition across the different deserts. This approach facilitated a comparative analysis of the migratory patterns prevalent within each desert group, allowing for a detailed understanding of migratory behavior in relation to desert ecosystems. Furthermore, the trophic niche categorization of the recorded species facilitated a detailed investigation into the trophic dynamics and ecological roles of avian species across various desert ecosystems.

To investigate the migratory behavior of avian species across selected deserts, categorized by trophic niches, a Sankey diagram was constructed in R software. The data in the Sankey diagram was represented by linking different deserts (source nodes) through their migration categories (first target nodes-M: Migratory, P: Partial Migratory, S: Sedentary) to their respective trophic niches (second target nodes). The width of the links were based on the count of avian species under each category.

Based on the AVONET dataset, a correlation matrix was established between the ten trophic niches to identify interrelation between the trophic niches of the selected deserts.

## 3. Results

Geospatial data curated from the Global Biodiversity Information Facility (GBIF) indicate that the ten major deserts analyzed harbor a combined total of 2,374 avian species. Notably, the Sahara Desert, being the largest, supports the highest number of species. Contrasts in species richness are apparent among deserts of similar sizes; for instance, the Arabian and Great Australian Deserts, as well as the Great Basin and Patagonian Deserts, and the Thar and Atacama Deserts do not exhibit comparable levels of species richness. On the contrary, statistical analysis reveals that deserts situated along the TCan exhibit significantly higher species richness, compared to those along the TCap, as illustrated in Figure 3.

**Figure 3:**
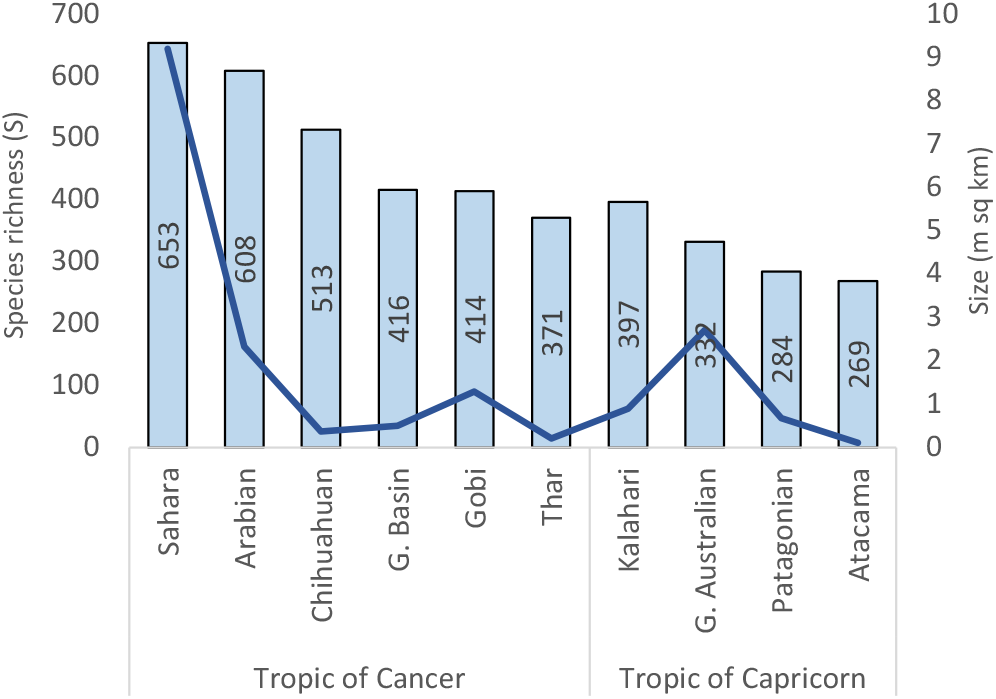
Variation in avian species richness across selected deserts. This figure presents avian species richness (S) relative to desert size (expressed in millions of square kilometers). Desert sizes are illustrated using a line plot on the secondary y-axis, highlighting that, with the exception of the Sahara, deserts along both the TCan and TCap are similarly sized. The primary y-axis features a bar plot displaying species richness, with specific values indicated at the center of each bar.

### 3.1. Variation in avian diversity across global desert ecosystems

The mean species richness calculated for deserts along the TCan was 464, in contrast to 295 for those along the TCap. Our analysis identified six species: *Ardea alba, Bubulcus ibis, Columba livia, Falco peregrinus, Passer domesticus*, and *Tyto alba*—as common across all deserts within TCap. Additionally, apart from these six species, the deserts under TCan harbored an additional 21 species that were common among them, resulting in a total of 27 shared species. Notably, *Tyto alba* was ubiquitous across all selected deserts included in the study, with the sole exception of the Gobi Desert. It was noted that distance did not play a crucial role in similarity in species richness between grouped deserts. For instance Sahara and Arabian deserts are closest (2293 km) of all the deserts and yet share only 9% (150) of their species. Great Basin and Chihuahuan on the other hand have greater distance and yet have 19% (299) common species.

In our analysis, the k-means clustering revealed that based on the avian species diversity, the 10 selected deserts form three clusters (Fig.4a). Group 1 was formed of Great Basin and Chihuahuan. Group 2 was formed of Sahara, Arabian, Thar and Gobi. Group 3 was formed of Kalahari, Great Australian, Patagonian and Atacama. It is important to note that, the Group 1 and 2 clusters when mapped to their location lie on the TCan, whereas the Group 3 are deserts that lie on the TCap (Fig. 1). Thus k-means clustering (Fig. 4a along with Figure 3 indicate the avian species richness is differentiated between TCan and TCap. In order to understand the type of correlation based on which the clusters were formed and their significance, we explored the correlation between the selected deserts. Although Fig.4a indicates, Sahara Arabian, Thar and Gobi as a single cluster in Group 1, the correlation matrix (Fig.4b) indicate a moderate positive correlation between all these four deserts. Notably, the Chihuahuan and Great Basin were the only deserts that displayed a significant positive correlation (r = 0.75). Conversely, though not significant, negative correlations were observed between the deserts of the group 3, i.e., those that lie in the TCap, except Patagonian and Atacama, which showed moderate positive correlation (r= 0.43) among themselves.

**Figure 4:**
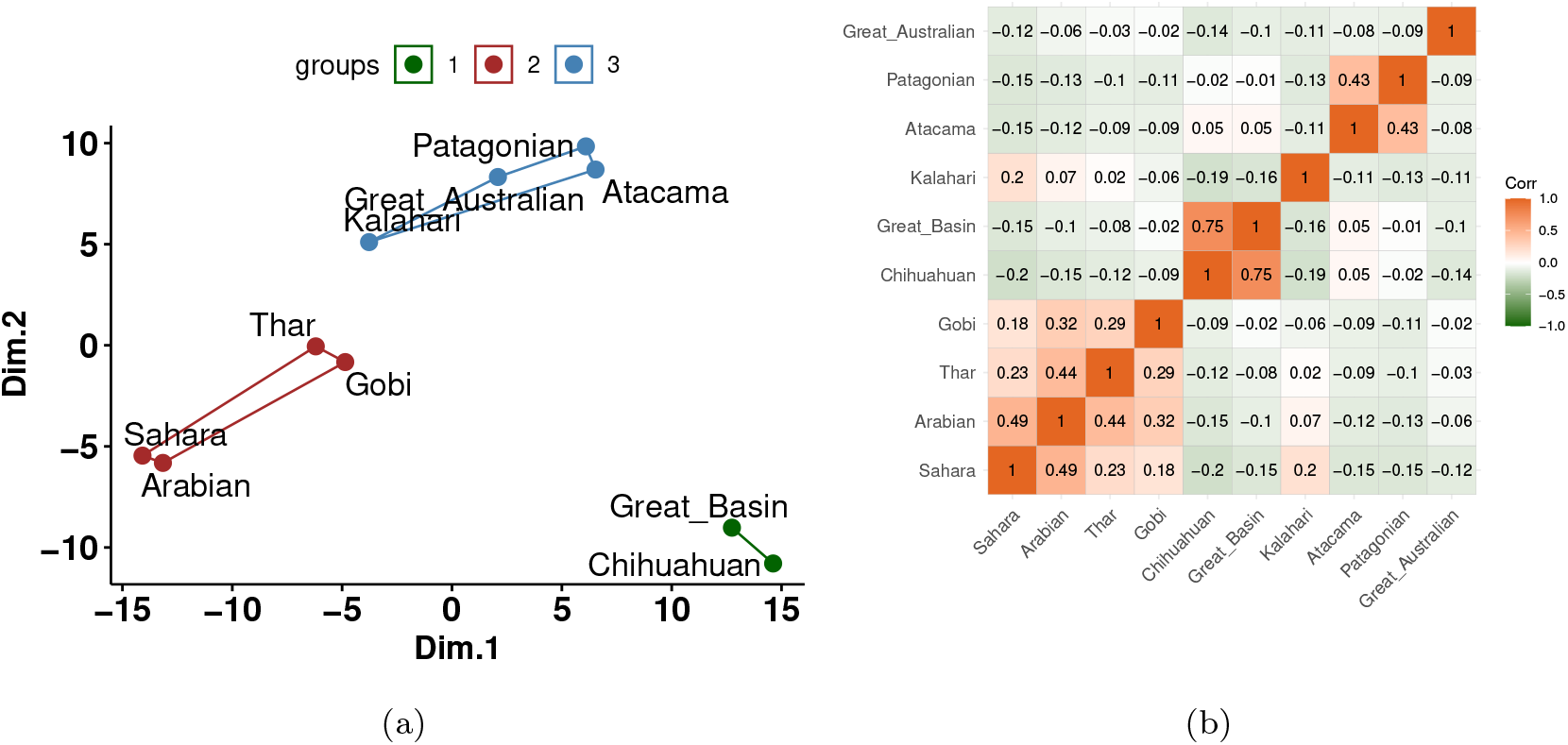
(a) Multidimensional Scaling (MDS) and k-means clustering with *k* = 3 resulted in three distinct clusters, represented by different colors: Group 1 in green, Group 2 in red, and Group 3 in blue. Geographically, Groups 1 and 2 are located in the Tropic of Cancer, corresponding to the Nearctic and Palaearctic realms respectively. Group 3 is situated in the Tropic of Capricorn, encompassing regions from the Australasian, Nearctic, and Neotropical realms. (b) A correlation plot (corrplot) based on a correlation matrix, with correlation coefficients (*r*) displayed at the center of each matrix cell. The correlation values range from -1 to 1, with values near -1 shown in green indicating a very high negative correlation in avian diversity, and values close to 1 in orange denoting a high similarity. The intensity of the colors signifies the degree of similarity or dissimilarity. The plot reveals five primary clusters based on positive correlations: the first cluster includes the Sahara, Arabian, Thar, and Gobi deserts; the second comprises the Great Basin and Chihuahuan deserts; the third contains the Patagonian and Atacama deserts; the fourth and fifth are the individual deserts of Kalahari and Great Australian, respectively, with Kalahari showing a moderate positive correlation with Palaearctic deserts, whereas the Great Australian shows no significant positive correlations.

### 3.2. Unique desert characteristics through functional traits and Migratory behaviour

The species richness data obtained from GBIF were mapped with the AVONET data and 1930 (80%) species could be matched to analyze the behavioural traits. Number of species under each category of migratory behaviour from the individual deserts are presented in Table (1). Analysis based on this dataset revealed persistent distinctions in species richness across these migratory categories between the two tropics; specifically, sedentary species richness was notably higher in TCap, while richness for both migratory and partial migratory categories was elevated in TCan. The disparity was most pronounced in migratory species, which exhibited significantly higher median values in TCan (Fig. 5a). These observations are corroborated by ANOVA outcomes, demonstrating statistically significant differences in avian species richness across the migratory categories and between the tropical zones, with respective p-values of 0.00635 and 0.00418 (Table 2). Additionally, the interaction effect between migratory category and geographic tropics, with a p-value of 0.00216, robustly indicates that the impact of migratory behavior on species counts is distinctly modulated by the tropical region.

**Table 1:** Migration category counts for Deserts in different Tropics.

| Tropics Deserts |  | Sedentary | Migratory | Partial migratory |
| --- | --- | --- | --- | --- |
| TCan | Sahara | 253 | 169 | 120 |
| TCan | Arabian | 166 | 214 | 121 |
| TCan | Thar | 110 | 117 | 80 |
| TCan | Gobi | 70 | 196 | 85 |
| TCan | Chihuahuan | 120 | 222 | 82 |
| TCan | Great Basin | 67 | 214 | 66 |
| TCap | Great Australian | 172 | 39 | 59 |
| TCap | Kalahari | 216 | 63 | 50 |
| TCap | Atacama | 107 | 58 | 49 |
| TCap | Patagonian | 107 | 54 | 67 |

**Table 2:** ANOVA results for Migration and Tropics.

| Source | Df | Sum Sq | Mean Sq | F value | Pr(>F) |
| --- | --- | --- | --- | --- | --- |
| Migration | 2 | 23138 | 11569 | 6.292 | 0.00635** |
| Tropics | 1 | 18422 | 18422 | 10.020 | 0.00418** |
| Migration:Tropics | 2 | 29463 | 14732 | 8.013 | 0.00216** |
| Residuals | 24 | 44125 | 1839 |  |  |
| **Significance codes:** 0 ‘***’ 0.001 ‘**’ 0.01 ‘*’ 0.05 ‘.’ 0.1 ‘ ’ 1 |  |  |  |  |  |

**Figure 5:**
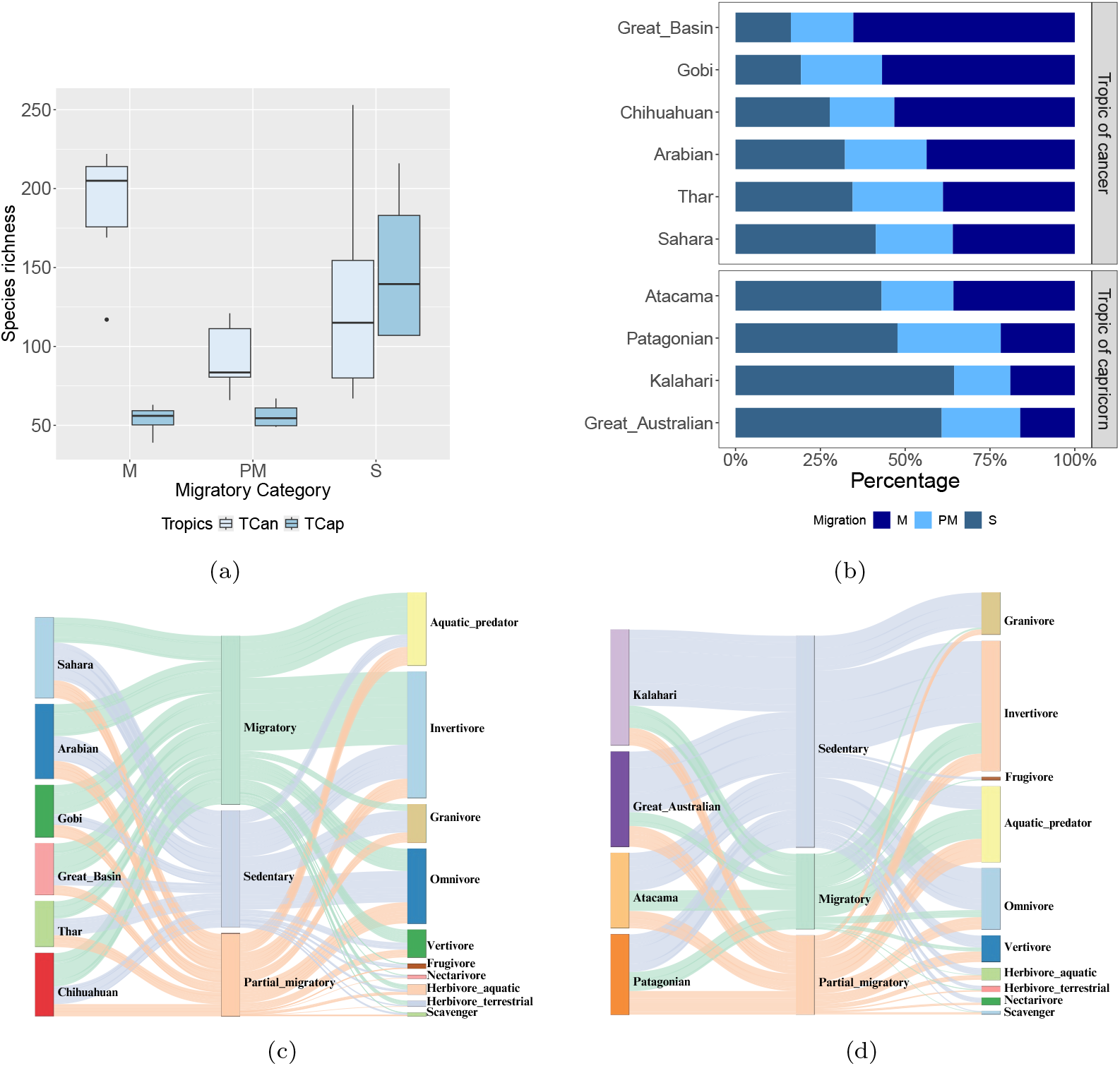
(a) Boxplot illustrating the variation in avian species counts across migratory categories (S = Sedentary, M = Migratory, PM = Partial Migratory) in different tropical zones (TCan and TCap). Notably, the migratory species tends to exhibit higher median counts in TCan, whereas in TCap, Sedentary and Partial migratory species show comparatively lower dispersion and median values. (b)Distribution of avian species in desert ecosystems categorized by migratory behavior. Migratory birds predominated in all TCan deserts, while sedentary species were more prevalent in TCap deserts. Partial migratory bird distribution exhibited consistency across all desert regions within both Tropics.(c) and (d) Trophic niche distribution of avian species across various deserts in Tcan and Tcap respectively, categorized by migratory behavior, based on data from Tobias et al. (2022). This Sankey diagram illustrates the flow of species from different deserts through their migration categories to their respective trophic niches. The nodes on the left (Sources) represent deserts belonging to each tropics and nodes on the right represents trophic niches of the birds. Each link represents the count of species transitioning from one category to another. The colors of the links correspond to the migration categories. Across all deserts studied, the composition of trophic niches is consistent, with Invertivores, Omnivores, and Aquatic predators comprising the largest proportions. More Invertivore species were migratory in TCan deserts but sedentary in TCap deserts. The trophic niches (Aquatic predators, Vertivores, and Granivores) that were predominant among migratory and partial migratory species in TCap also predominated among sedentary species in TCan.

According to the data depicted in Figure 5b, the deserts of TCan showed a higher percentage of migratory species, ranging from 35% to 65%. In contrast, sedentary birds were more prevalent in the deserts of TCap, constituting between 43% and 64% of the species (Fig. 5b). The frequency of partial migratory birds was relatively stable across all deserts within both TCan and TCap, with figures varying from 18% to 30%. This analysis highlights how migratory patterns might influence the observed correlation patterns between these distinct geographic zones. It is also important to note that among the deserts of TCan, the Nearctic deserts (Great Basin and Chihuahuan) inhabited higher number of migratory species, when compared to the Palaearctic deserts (Arabian, Thar and Sahara). Among the deserts of TCap, the Neotropical deserts (Atacama and Patagonian) inhabited higher number of migratory species when compared to Afrotropical and Australasian deserts.

### 3.3. Impact of Trophic Niche Structures on Migratory Bird Patterns in Desert Environments

To investigate the underlying causes of observed differences in migratory behavior between avian species across deserts in two tropics, trophic niche data from AVONET were employed. Figure 5c demonstrates that Invertivores are the predominant trophic niche while Scavengers are the least dominant across the selected deserts of Tcan. The second target nodes at the right end of the diagram distinguished between the most dominant trophic niches (Invertivores, Aquatic predators, and Omnivores) and the least dominant ones (Granivores (G), Vertivores (V), Herbivore aquatic (Ha), Herbivore terrestrial (Ht), Nectarivore (N), Frugivore (F), and Scavengers (S)). The links based on their width effectively classified the deserts based on avian migratory behavior, highlighting distinct trophic niche selections across the regions. The green links indicated that the migratory species in all deserts under the TCan designation were primarily Invertivores, succeeded by Aquatic predators and Omnivores. Conversely, the blue and peach coloured links revealed that these same niches were prevalent among the sedentary and partial migratory species of TCap deserts (Fig. 5d). The peach links comprised partial migratory birds show a wider spectrum of trophic niches used and no predominance to any particular trophic niche. Additionally, it was noted that Omnivores were less common among the migratory species of TCap deserts compared to those of TCan deserts.

## 4. Discussion

Understanding the intricacies of avian biodiversity across the world’s deserts necessitates an integrated approach that considers geographic, climatic and ecological factors. We utilized crowd-sourced eBird data and comprehensive ecological datasets to analyze the biodiversity of ten major deserts, covering 12% of the Earth’s land area (149.6 million km^2^) and supporting 22% of global avian species. Our study reveals significant variability in species richness among these deserts (Fig. 3), challenging the notion that desert size is a primary determinant of biodiversity (Levin, 2013; Sorte et al., 2022; Lomolino and Rosenzweig, 1996).

Despite similarities in geographic climatic conditions (Coelho et al., 2023) like aridity, temperature, and precipitation, the deserts exhibited only six common bird species, underscoring their unique ecological profiles. Our findings indicate that geographic location specific ecological conditions (Fig. 5c and 5d), rather than desert size (Fig. 3), drive differences in avian species richness. Deserts near the TCan were notably more species-rich than those near the TCap, reflecting the complex interplay of climatic and geographic factors (Terborgh, 1973).

Intriguingly, TCan deserts host a higher percentage of migratory species compared to TCap deserts (Table 2, Fig. 5a and 5b), suggesting that migratory behavior significantly influences overall species richness. While both regions fall within the same latitudinal diversity gradient, TCan deserts benefit from a greater influx of migratory species due to the wide range of trophic niches occupied by sedentary species (Fig. 5c and 5d). This reduces competition for migratory species and supports higher overall richness due to increased environmental heterogeneity (Fargallo et al., 2022) in TCan.

Unlike other ecosystems where vegetation productivity is a key driver of species richness (Wright, 1983; Hobi et al., 2021), desert biomes rely heavily on detritivores for nutrient and energy cycling (Crawford, 1979). This unique dynamic is corroborated in the correlation between migratory populations and species richness, particularly among invertivores (Fig. 5c and 5d), signifying invertebrates are crucial in determining avian diversity in deserts across the globe.

Biogeographic realms, traditionally defined by species distribution patterns (Russel, 1876), provide another layer of insight. Our analysis places the studied deserts within seven of Holt’s redefined realms Holt et al. (2013), revealing three primary clusters based on avian diversity: Nearctic, Palaearctic, and a combined Afrotropical, Neotropical, and Australasian realm. This clustering contrasts with the correlation matrix (Fig. 4b), which aligns deserts with five biogeographic realms, suggesting that avian species richness is more closely linked to tropical locations than to realm classifications.

The implications of our findings are significant for conservation efforts. Ecosystems with higher species richness are generally more resilient to disturbances, including climate change (Isbell et al., 2015; Tilman and Downing, 1994). Our results suggest that TCan deserts, with their diverse trophic niches and higher species richness, are likely to be more resilient compared to TCap deserts. Conversely, TCap deserts, characterized by lower species richness and narrower trophic niches, may face increased vulnerability to environmental changes and higher extinction rates (Emerson and Kolm, 2005; Weeks et al., 2022).

In light of these findings, it is imperative to prioritize conservation strategies that enhance the resilience of desert ecosystems. Protecting regions with high avian diversity, particularly those in the TCan deserts, and mitigating anthropogenic stressors in more vulnerable areas like TCap deserts are essential measures. This study’s scope was limited to two ecological characteristics: migration and trophic niches. A more comprehensive analysis incorporating additional ecological traits could elucidate further patterns in desert biodiversity,their diverse adaptive mechanisms and underlying causes of these differences. Moreover, future research should investigate the ecological dynamics of other taxonomic groups to ascertain whether their conservation requirements are congruent with those of avian species.

## 5. Conclusion

This study provides the first comprehensive analysis of avian biodiversity and resilience in desert ecosystems across the Tropics-TCan and TCap. Our findings reveal that TCan deserts exhibit higher species richness, particularly among migratory birds, and greater resilience to climate change compared to TCap deserts. The adaptability of sedentary species to a broader trophic spectrum in TCan deserts reduces niche competition, facilitating a higher influx of migratory species. We recommend targeted conservation strategies to protect the unique avian diversity in TCan deserts and mitigate extinction risks in TCap deserts.

## Authorship contribution statement

**Manasi Mukherjee** contributed in conceptualization, data curation, investigation, methodology, analysis, writing and reviewing. **Mitali Mukerji** contributed to supervision, writing, reviewing and editing.

## Data Availability Statement

All data generated or analysed during this study are included in this published article (and its Supplementary Information files).

